# Differences in density: taxonomic but not functional diversity in seaweed microbiomes affected by an earthquake

**DOI:** 10.1101/2023.02.08.527737

**Authors:** William S. Pearman, Sergio E. Morales, Felix Vaux, Neil J. Gemmell, Ceridwen I. Fraser

## Abstract

Host-associated microbial communities can make important contributions to host health, and are shaped by a range of different factors ranging from host condition, environmental conditions, and other microbes. Disentangling the roles of these factors can be particularly difficult as many variables are correlated. Here, we leveraged earthquake-induced changes in host density to identify the influence of host density on microbiome composition. A large (7.8 magnitude) earthquake in New Zealand in 2016 led to widespread coastal uplift of up to ~6m, sufficient to locally extirpate some intertidal kelp populations. These uplifted populations are slowly recovering, but intertidal kelps remain at much lower densities than at nearby, less uplifted sites. By comparing the microbiome of the low and high density sites using 16S amplicon sequencing, we observed that low density populations had higher beta-diversity than high density populations with regards to taxonomic variability, while no beta-diversity differences were observed between functional categories. Using phylogenetic and taxonomic turnover approaches, we determined that dispersal limitation shapes low density populations to a greater extent, while homogeneous selection shapes high density populations to a greater extent. Our findings shed light on microbiome assembly processes, particularly highlighting that large-scale disturbances that affect host density can dramatically influence microbiome structure.

## Introduction

Large-scale disturbances can create natural experiments which can enable examination of how a range of ecological factors affect microbial communities (1). Disturbances can dramatically alter the environmental, geological, or environmental landscape and thus present ideal systems to understand drivers of microbiome assembly and composition. Volcanic activity leading to the creation of new islands has enabled the study of community assembly from the earliest stages, and even geological activity occurring millions of years ago continues to provide insights towards the evolution and ecology of host-associated microbial communities (1,2). On shorter time scales, natural disasters that create widespread ecological disturbance can provide direct insight into microbial community dynamics (3,4) in more realistic ways than tightly controlled laboratory experiments that often fail to capture the complexity of natural environments.

The population density of a host species has the potential to constrain microbiome diversity due to interactions among hosts resulting in newly recruited host individuals obtaining microbes via horizontal transmission (5). In essence, increased host density can reduce dispersal limitation of microbes. Clear understanding of this potential relationship is, however, complicated by a range of confounding variables because population density or size is often related to the environmental suitability of a habitat (6). This relationship can make it difficult to decouple the contribution of host density to the microbiome from the contribution of the environment. Furthermore, most research to date has largely focused on experimental approaches to these questions, which are removed from the ecological context where they would normally occur (1). Natural disasters such as marine heatwaves and earthquakes can lead to long-term changes in host density, well after the environmental influence of the disaster has abated, enabling decoupling of environmental conditions from population density.

Previous research has found that environmental conditions, host anatomy (7), epiphytes (8), and disease (9) strongly influence the microbiome structure of marine macroalgae. Emerging research also indicates a strong contribution of the microbiome towards macroalgal health (9) and growth (10), and further evidence suggests that these microbiomes tend to be relatively resistant to change (11). To complicate this picture, host genetic constraints may limit flexibility in microbial invasion (12), highlighting the complexity of microbiome assembly processes in marine macroalgae.

Here, we use the opportunity presented by the 7.8 magnitude 2016 Kaikōura earthquake in Aotearoa New Zealand to examine the influence of host density on microbiome composition of southern bull kelp, *Durvillaea antarctica*. This earthquake led to non-uniform coastal uplift of up to ~6 m (13,14), leading to localized extirpation of some southern bull kelp populations (15). The patchiness of the uplift resulted in some affected ecosystems being geographically close to relatively unaffected populations, and we leveraged this proximity to undertake a comparative study to understand the influence of tissue type, host age, and population density on the microbiome.

Since the 2016 earthquake, southern bull kelp has been recolonizing coastlines from which it was extirpated. This recolonization process has been quite slow, and although adult hosts now occur on these coastlines once again, they often occur at remarkably low abundances (15). While population densities are known to vary among *Durvillaea* species (16), habitat cover for intertidal *Durvillaea* has been found to be substantially lower at locations more severely uplifted by the 2016 earthquake (17). Dispersal to dramatically uplifted sites (>2 m) from which *Durvillaea* was extirpated would have most likely occurred via the rafting of adults that release eggs and sperm to form zygote settlers (15,18), and by the time these young had become established, their parents – the adult dispersers – would in most cases have died or drifted away. The first southern bull kelp to settle in such locations therefore established themselves in the absence of other intertidal kelp, and thus had to obtain their microbiota without much, if any, horizontal transmission from conspecifics. In contrast, at less uplifted sites nearby (<2 m uplift), density crashes were not so severe and population recovery has been much swifter. We hypothesised that at the most uplifted sites, where populations were extirpated, kelp microbiomes would be distinguishable from those of kelp at less uplifted sites nearby, because of the need for the former to re-establish microbiomes without transfer from dense stands of conspecifics. This elegant natural system allows us to directly examine not only the microbial differences between extant and re-colonizing populations, but also the ecological drivers behind these differences.

## Methods

In November 2020, we collected paired microbiome and tissue samples from *D. antarctica* ‘NZ North’ lineage (as described in (19)) individuals at four sites in the north Canterbury region of New Zealand (Fig. 1). These sites were chosen in pairs, where one site in each pair experienced significant geological upheaval in the 2016 earthquake, and the other site did not. The first pair contained southern locations: Kaikōura (0.8 m uplift; high density) and Waipapa Bay (4.4 m; low density), whereas the second contained the northern locations: Cape Campbell (0.5 m; high density) and Ward Beach (2.2 m; low density).

**Figure 1.**
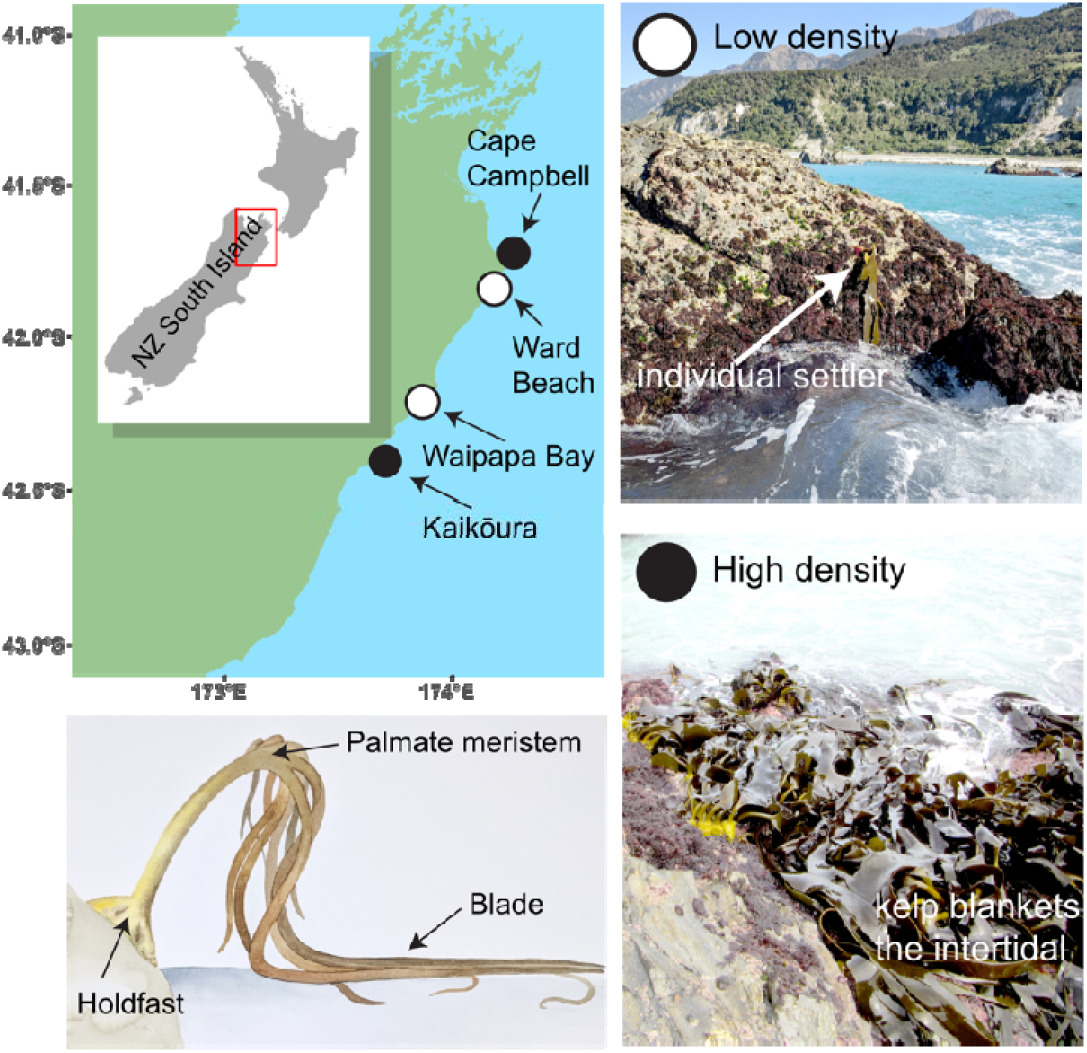
Sampling sites in North Canterbury, New Zealand. Points in white are sites which experienced significant uplift and are low density populations, whereas black points are sites which did not experience significant earthquake induced density shifts (example photos on right). The sampled parts of the host (*Durvillaea antarctica*) are indicated (bottom left panel).

At each site, we sampled 18 hosts, comprising 12 mature adults and 6 juveniles, however due to extremely low density at some sites fewer samples were collected (Table 2). From each individual, we collected a host tissue clipping that was preserved via desiccation with silica gel, and triplicate microbiome samples via surface swabbing. Two of these microbiome samples were collected from different blades, 30 cm from the tip, and one sample from the palmate meristem region (see Figure 1) of the kelp; all were collected by swabbing back and forth over a 5 x 5 cm area with a Qiagen OmniSwab. Swabs were stored in sterile DESS (DMSO EDTA Saturated Salt solution) immediately upon collection and were stored at −20°C while in the field, then transferred to a −80°C freezer on return to the laboratory. Four substrate swabs were collected per site using the same methodology on rock surfaces adjacent to kelp holdfasts.

At each site, we also collected four 2L water samples from close to the kelp, processed using a bleach-sterilized vacuum filter, with 0.22uM polycarbonate filters that were subsequently stored in DESS and frozen.

### DNA Extraction

DNA was extracted from all samples using the Qiagen PowerSoil Kit, following manufacturer’s instructions (Supp. Methods) with all optional steps, and with 2 rounds of bead beating at 25Hz for 5 minutes per round using a Domel MillMix. DNA was eluted in nuclease free water, using a two-step elution process of 50uL of water in each step, with a 5 minute incubation at room temperature between elutions. DNA was then transferred to a 96 well microtiter plate and dried using an Eppendorf SpeedVac.

### Library Preparation

16S library preparation was carried out at Argonne National Laboratories (ANL), following standard earth microbiome project sequencing protocols (20,21). In short, DNA was amplified using updated 515F (5’: GTGYCAGCMGCCGCGGTAA) and 806R (5’: GGACTACNVGGGTWTCTAAT) primer pair to amplify the V4-V5 region of the 16S rRNA gene. Amplicons were then pooled equimolarly at ANL and sequenced on an Illumina MiSeq with a 2 x 150 bp library to produce amplicons with an approximately 50 bp overlap.

### Data Processing

16S amplicon reads were quality filtered and processed in DADA2 (v. 1.26) (22), removing PhiX reads, and truncating reads at the first instance of a base with quality less than two. Chimeric reads were removed using consensus identification in DADA2. The resulting ASV set was then decontaminated using the *decontam* (v. 1.16) R package (23) with a significance threshold of 0.1. Finally, ASVs were rarefied to a depth of 3,000 reads per sample using the rarefy_even_depth function in the R package *phyloseq* (v. 1.4) (24). Phylogenetic relationships between ASVs, and taxonomic classifications of ASVs, were performed by employing SATe-enabled phylogenetic placement (SEPP) (25,26) to graft ASVs onto a pregenerated GreenGenes 16S database. To assign ASVs to taxa, the nearest tip that was not another ASV in the phylogeny was identified and treated as the taxon.

### Analysis

Richness estimates were calculated using the estimate_richness function in *phyloseq*, and ordination was performed using Bray-Curtis distance with the ordinate function in *phyloseq*, with a k of 3. Beta-diversity was calculated using the betadiver function in *vegan*, following the method of Whittaker (1960). Differences in beta-diversity were calculated based on distance to centroid, rather than median, alongside a bias adjustment for small sample sizes as implemented by the *betadisp* function in the *vegan* (v. 2.6.2) R package. Following betadiversity tests, comparisons with non-significant differences in beta-diversity were tested for differences using PERMANOVA as implemented by the adonis2 function in *vegan* (27).

Functional annotation of ASVs was conducted using FAPROTAX v 1.2.6 (28) based on the SEPP informed taxonomic assignments. Core microbiome, both functional and taxonomic, was identified by using the core_members function in the R package *microbiome* (v. 1.18) (29), on a relative abundance transformed ASV table, using a threshold of 80 % (i.e., microbes in >80 % of samples were considered ‘core’). All R analyses were conducted using R 4.1.1 (30).

Because the functional analyses collapsed all ASVs into 92 functional groups, we conducted a power analysis to ascertain if a dataset with 92 groups would be capable to detect a small effect size of 0.1. This was done with the *wmwpow* (v. 0.1.3) R package (31) and revealed that this dataset, with at least 85 samples per group would be sufficient to detect such a small effect size. Therefore, we proceeded with three analyses; comparing functional beta-diversity (calculated as above following Whittaker 1960) based on 92 groups, to ASV beta-diversity calculated from both a sub-sample of 92 ASVs, and the full complement of ASVs. These analyses were done by conducting Mann-Whitney U tests in the *coin* (v. 1.4-2) R package (32), and using the observed Z value to calculate effect sizes by dividing *Z* by the square root of the number of samples.

To understand the relationship between community structure and ecological processes, we employed the iCAMP framework developed by Stegen et al. (33) and extended by Ning et al. (34), using the RC.cm and BNTI.cm functions in the R package *iCAMP* (v. 1.5.12) (35). These functions calculated pairwise Raup-Crick (RC) and Beta-Nearest Taxon Index (βNTI) based on pairwise comparison of samples. Categorization of βNTI and RC into ecological processes is outlined in table 1. Homogeneous selection occurs when selective pressures are constant and low phylogenetic turnover is expected, while heterogeneous selection occurs when selective pressures are variable leading to greater than expected phylogenetic turnover. After selection has been accounted for (values of |βNTI| ≤ 2), the remaining community structure can be explained by dispersal and drift. Dispersal limitation occurs when taxonomic turnover is higher than expected (RC > 0.95), while homogeneous dispersal occurs when taxonomic turnover is lower than expected (RC < −0.95). Remaining community structure can then be attributed to ecological drift (variation in community structure arising from chance).

**Table 1.**
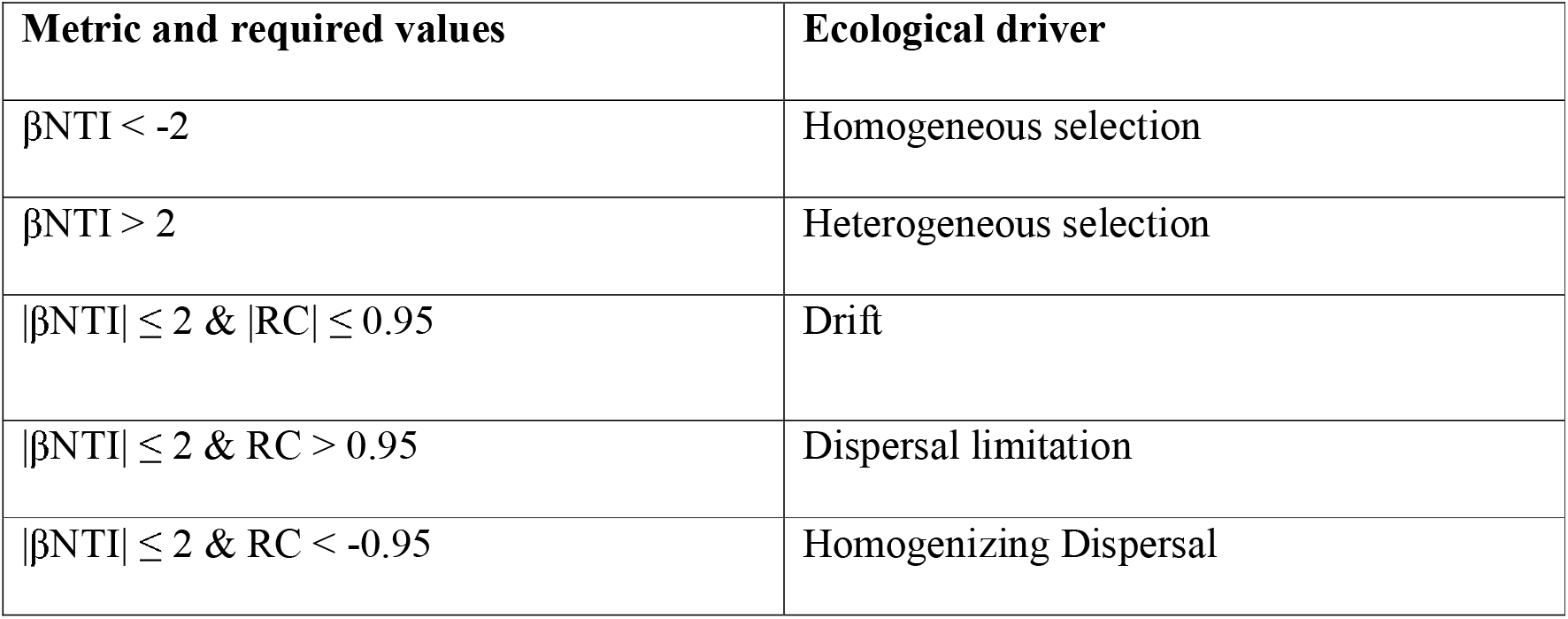
Breakdown of metrics and their relationship to different drivers of community structure. βNTI refers to Beta Nearest Taxon Index, while RC refers to the Raup-Crick index.

## Results

### Sequencing summary

A total of 235 microbial samples were sequenced successfully (Table 2), with an average of 34,070 merged reads retained per sample following DADA2 quality filtering. Following decontamination and rarefaction, 5,754 ASVs were retained for further analyses.

**Table 2.** The number of successfully sequenced samples for each location and sample type.

|  | Cape Campbell<br>(high density) | Kaikōura<br>(high density) | Waipapa Bay<br>(low density) | Ward Beach<br>(low density) |
| --- | --- | --- | --- | --- |
| Substrate | 4 | 4 | 3 | 4 |
| Seawater | 4 | 4 | 4 | 4 |
| Palmate Meristem (Adult) | 12 | 12 | 11 | 11 |
| Blade (Adult) | 24 | 24 | 22 | 22 |
| Palmate Meristem (Juvenile) | 5 | 7 | 4 | 6 |
| Blade (Juvenile) | 13 | 12 | 8 | 12 |

#### Microbial communities are distinct between environments

We found that seawater and substrate associated microbiomes had 3-4 times more ASVs than seaweed microbiomes (356.2 and 281.5 ASVs on average for seawater and substrate, compared to 80 ASVs on average for kelp, Kruskal-Wallis test – Substrate vs Kelp P= 2.6×10^-12^, Seawater vs Kelp=1. 6×10^-10^) (Figure 2). Furthermore, meristematic tissue tended to have lower ASV richness than blade tissue (61.1 vs 97 on average, Kruskal-Wallis test P = 6.5×10^-5^) (Figure 2). However, no difference in richness between age classes was observed (Kruskal-Wallis test P = 0.962).

**Figure 2.**
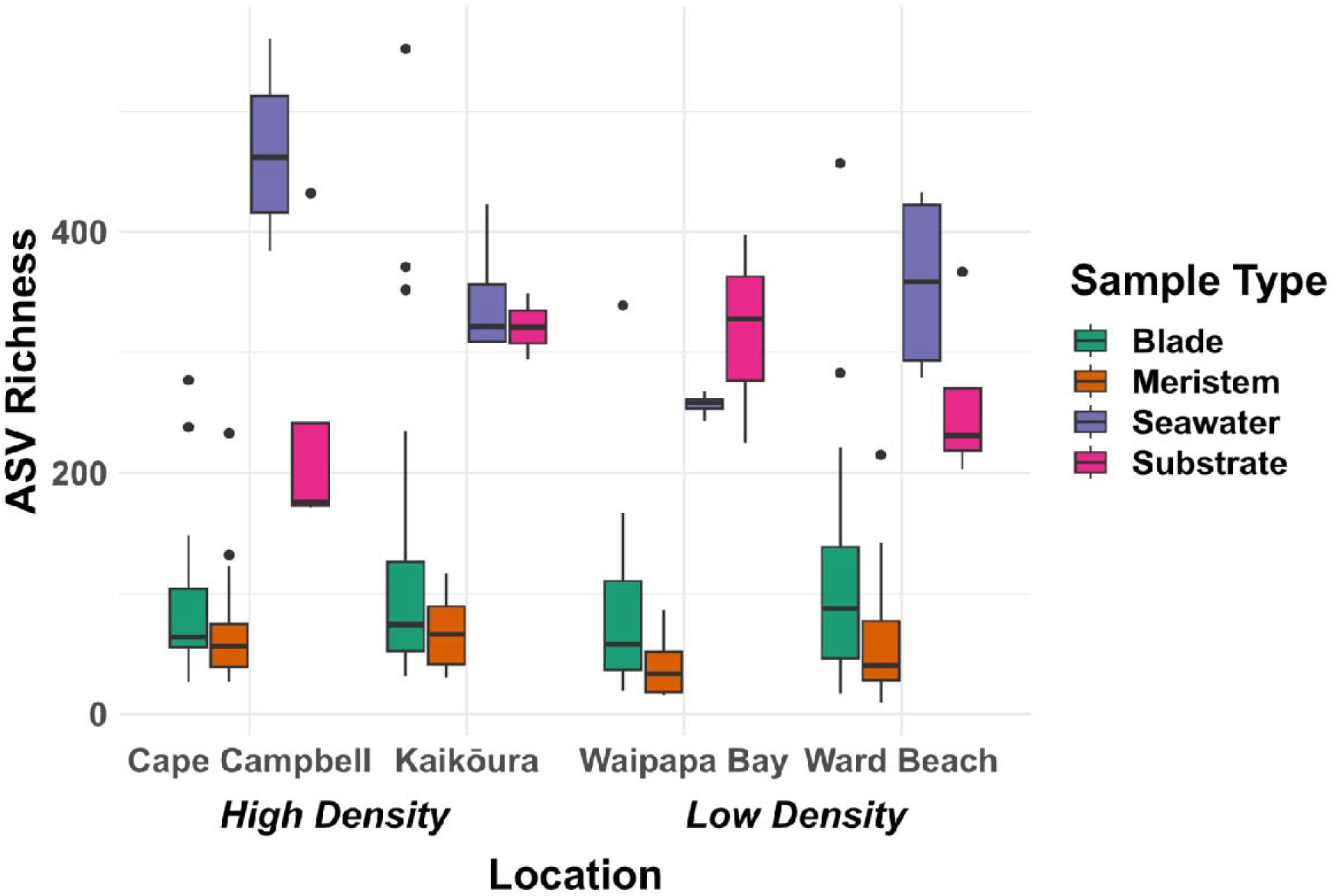
Amplicon Sequence Variant (ASV) richness based on rarefied data, for different sample types and locations.

Multivariate analysis of homogeneity of variance using PERMDISP revealed that low density populations had higher beta diversity than high density populations regardless of tissue type, while no differences in beta-diversity were observed when examining seawater and substrate communities (Table 3). While differences were present between density classes, none were observed between tissue types or age classes. Additionally, PERMANOVA analyses also revealed that within density classes, locations show significant differences (Table 3).

**Table 3.**
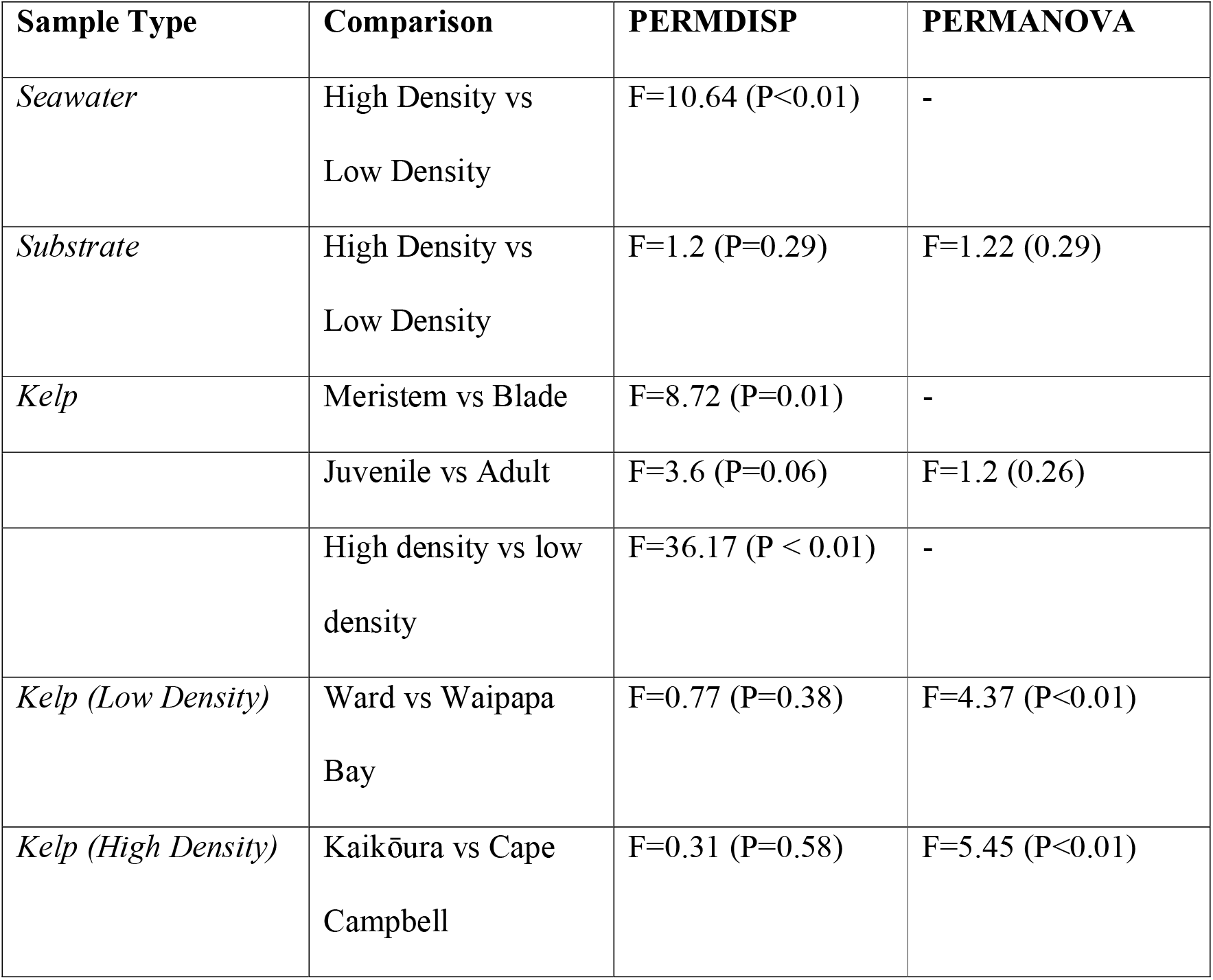
PERMDISP and PERMANOVA results for differing microbial community comparisons. Absent PERMANOVA values are present due to significant differences in beta-diversity rendering PERMANOVA inappropriate.

Analysis of average distance to centroids showed that low density populations had higher distance to centroids, further supporting the observation of increased beta-diversity associated with low density populations (Mann-Whitney U test, W=6947, p<0.001, Figure 3). While there were significant differences between locations within density groups, low density populations consistently had higher beta-diversity than their high-density counterparts. Furthermore, blade tissue had higher beta-diversity than meristem tissue and trended towards having higher beta-diversity than adults.

**Figure 3A:**
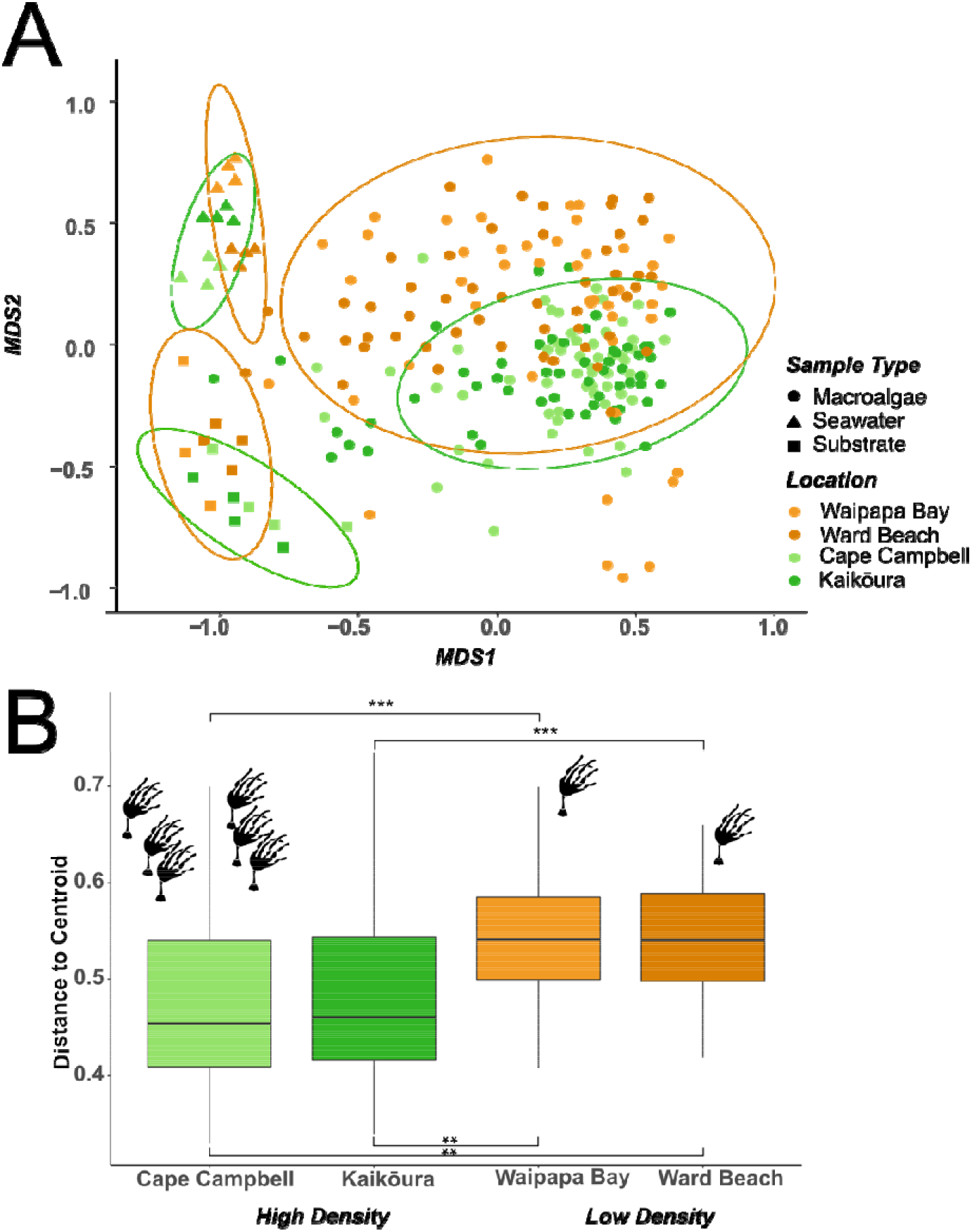
Non-metric Dimensionality Scaling plot of microbial communities in low density (shades of brown) and high density (shades of green) locations. Ellipses represent clustering at the density class level. Stress level for this NMDS is 0.19. 3B: Distance of individual communities to the centroid of each community comparison. Centroids are based on Principal Coordinate Analyses using Sorensen dissimilarity based on Whittaker (1960). Significance values are based on Mann-Whitney tests, non-significant comparisons are not displayed

Non-scaled dimensionality reduction (NMDS) approaches revealed distinct clusters of seawater, substrate, and macroalgae associated microbial communities. The previously observed differences in beta-diversity were also shown on the NMDS, with greater variation in low density communities relative to high density communities (Fig. 3).

#### Microbiomes are functionally, but not taxonomically, conserved across host density levels

Using FAPROTAX, we assigned ASVs to functional groups, then we identified the core taxa and functions in the *D. antarctica* microbiome. We found that for low density sites there were 2 core ASVs, assigned to *Granulosococcus* and *Gammaproteobacteria*. In contrast, samples from high density populations possessed 11 core ASVs assigned to *Flavobacteriales*, *Leucothrix sargassi, Saprospiraceae, Acidimicrobiales*, and *Flavobacteriales* in addition to *Gammaproteobacteria* and *Granulosicoccus*. Both density classes included chemoheterotrophic and aerobic chemoheterotrophic functional groups, however high density samples also possessed intracellular parasites as a core functional group.

We then conducted beta-diversity and PERMANOVA analyses based on the functional groups identified in the kelp microbiomes rather than the ASVs. We observed no differences between density groups with regards to functional beta-diversity (Z=-1.68, p = 0.094, r=0.11). Equally powered tests at the ASV level revealed significant differences between density groups (Z=-4.12, p<0.001, r=0.3), this result was similar in effect size to the analyses conducted with the full complement of ASVs (Z=-5.01, p<0.001, r=0.35). Finally, there were no significant differences between density classes within pairs of populations based on PERMANOVA (F=2.76, p = 0.071).

### Ecological processes

Finally, to determine what underlying ecological processes were shaping community structure in *D. antarctica* microbiomes, we applied the framework developed by Stegen et al (33), and Ning et al (34). This framework examines the degree of taxonomic and phylogenetic turnover in communities relative to a null model to identify the roles of selection, dispersal, and drift. Using the beta nearest taxon index (βNTI), we found that homogenous selection accounted for 71.5% of community structure in high density populations, and 67.4% of community structure in low density populations (Fig. 3; Table 4). Additionally, we observed that dispersal limitation explained 1.7% of community structure in low density populations, and 0.2% in high density populations; other metrics were qualitatively similar between high and low density populations (Table 4).

**Table 4.** Percentage contribution of different ecological factors to community assembly for Durvillaea antarctica microbiomes under different host densities. See table 1 for definitions of ecosystem drivers.

|  | Homogeneous Selection | Heterogeneous selection | Drift | Limiting dispersal | Homogenizing dispersal |
| --- | --- | --- | --- | --- | --- |
| High density | 71.5% | 0% | 28% | 0.2% | 0.3% |
| Low density | 67.4% | 1.1% | 29.3% | 1.7% | 0.5% |

Furthermore, we found that in high density populations microbial communities were skewed towards lower Raup-Crick values, suggesting greater homogeneity of microbiome structure within these communities, while low density populations had a more even distribution of Raup-Crick values (Figure 4).

**Figure 4.**
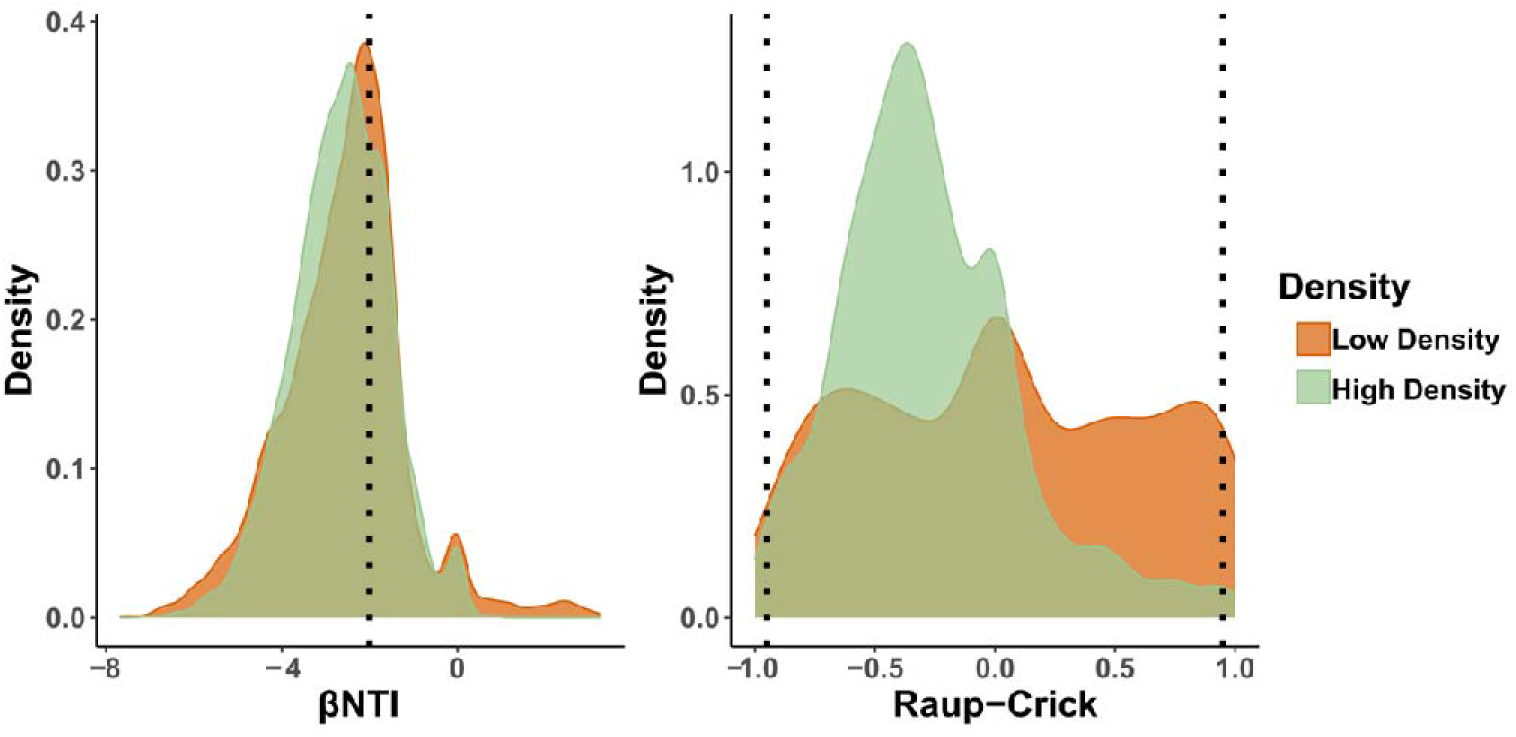
Distributions of β-nearest Taxon and Raup-Crick Indices for pairwise community comparisons within low (red) and high (blue) density communities. Dashed lines indicate which community comparisons are dominated by different ecological drivers. Values of βNTI < −2 indicate dominance of homogenous selection. Values of Raup-Crick < −0.95 indicate dominance of homogenizing dispersal and values of Raup-Crick > 0.95 indicate dominance of dispersal limitation in community assembly.

## Discussion

Our results support our hypothesis that host density can influence microbiome assembly, with *D. antarctica* microbiomes showing greater beta-diversity in low density populations that experienced significant disturbance from severe uplift (>2 m), compared to high density populations that received less disturbance from moderate uplift (<2 m). Interestingly, these differences were only identified at the ASV (taxonomic) level rather than for functional categories, suggesting that functional redundancy may compensate for differences in ASV composition (36).

Our finding that microbial communities associated with both the substrate and surrounding seawater were distinct from kelp microbiomes aligns with observations from other studies on macroalgae (11,37). Many of the core ASVs associated with *D. antarctica* belong to groups previously known to have macroalgal associations, such as *Granulosicoccus* and *Leucothrix* (7,8,38). Further examination of core microbial functions associated with *D. antarctica* revealed enrichment of heterotrophic bacteria in all populations. These results are also echoed by recent metagenomic work examining the microbiome of *Nereocystis luetkeana*, which are dominated by microbes associated with heterotrophy, nitrate reduction, and phototrophy (39). While few differences were observed between density classes with regards to function, the increased prevalence of intracellular parasites in high density populations may be the result of greater physical interactions between hosts which has the potential to promote the spread of parasites and diseases (40), whereas in low density populations there is reduced spread of diseases (41).

Consistent with previous research on other taxa (7,42), we found that microbiome composition differed between tissue types, with tissue from the palmate meristem being of lower richness than blade surfaces. These differences might be driven by two primary factors: differential exposure to environmental conditions, and different tissue associated growth rates (43,44). First, *D. antarctica* blades have high exposure to the marine environment relative to the palmate meristem region due to their buoyancy and length resulting in extension into the water column (in contrast, the palmate region is often out of the water at low tide: Figure 1). Secondly, palmate tissue tends to more resistant to hydrodynamic stress than blade tissue and thus is also likely to experience reduced erosion or breakage than the blade (44–46). As a result, we would anticipate that blade-associated microbial communities would have greater beta-diversity than meristems due their greater environmental exposure and probability of tissue erosion. Finally, meristematic tissue tends to be younger and under active growth (47) and is frequently enriched for growth related hormones which may lead to additional selective pressures (48) on the meristem resulting in a more homogeneous meristematic microbiome relative to the blade.

Our finding that host age class did not affect microbiome structure suggests that both recently recruited kelp and mature kelp have extremely similar microbial communities. Given the potential for collinearity between population density and host age, the absence of age-related microbiome structure supports our inference that it is population density that drives microbiome beta-diversity in *D. antarctica*.

Interaction among hosts is likely to underpin microbiome assembly dynamics. Population size has been found to affect microbiome beta-diversity in pikas (small rodent-like mammals), attributed to increased social interaction in larger populations (5). Because *D. antarctica* is a non-mobile species social interaction can be discounted; nevertheless, dense populations of *D. antarctica* will have greater physical interaction between neighbouring individuals. In low density populations, most kelp occurred individually, whereas in high density populations, kelp occurred in dense clusters where individuals frequently touch each other. The absence of direct physical interaction for many individuals in low density populations means that those microbiomes are more likely to be dispersal limited and thus undergo succession processes (and perhaps, to some extent, recruitment) independently (49,50). In essence, if we view kelp populations as microbial metapopulations (51), then low density kelp populations have lower connectivity and thus each microbiome undertakes its own independent successional trajectory – resulting in greater beta diversity.

Disregarding the potential of physical interactions, violent wave action is characteristic of the environment *D. antarctica* occurs in, and thus we suggest that this wave action may have an erosion-like effect on the kelp biofilm (52). Biofilm erosion would likely enrich the seawater for kelp-associated organic material, including eroded microbes, and thus the wave action may alter the seawater microbial composition, and enrich it for kelp-associated microbes (48). This effect would be stronger in high density populations, and thus we would expect that increased population density would reduce the independence of individual kelp microbial succession trajectories.

These hypotheses are further supported by the differing contributions of ecological processes to microbiome structure, as identified using the iCAMP framework. Broadly, we observed that like other marine microbiomes (53), both high and low density populations were dominated by homogenous selection suggesting that relatively constant selection and similar selective pressures influence microbiome assembly regardless of host density. However, drift and dispersal limitation were greater contributors to microbiome structure in low density populations than for high density populations. Independence of microbiome successional trajectories is anticipated to increase the influence of drift, because interactions between hosts promote microbial dispersal and lessen the probability that a microbial taxon is lost (50,54,55). For the same reason, we would also anticipate dispersal limitation to be a greater factor in low density populations, as frequency of physical interactions may shape the microbiome of *D. antarctica* as they do in other species (56). Furthermore, many sampled hosts will be recent recruits (post-earthquake – most adults probably just one or two years old) in low density populations and thus their microbes will have had fewer dispersal opportunities than for high density (older) kelp, because of time limitations. Together the fewer dispersal opportunities both temporally and because of reduced physical interactions may explain the increased influence of drift and dispersal limitation on the microbiome of *D. antarctica*.

Our finding that functional groups did not differ much with population density suggests that despite taxonomic differences, there is considerable functional redundancy in the kelp microbiome, similar to observations within sea lettuce (*Ulva;* (57,58). Functional redundancy in macroalgal microbiomes may be particularly important in the case of population establishment when ‘typical’ kelp-associated microbes may not be available (12). Our results suggest that *D. antarctica* is able to host a flexible assortment of microbes that fulfil the same core functions. These results echo findings from other research indicating that microbiome flexibility could facilitate species invasions (12,59), especially because the recolonization process of *Durvillaea* following the earthquake is analogous to species invasion dynamics (see other examples; (60). This work aligns with the generalist host hypothesis which posits that hosts which are more flexible in terms of their microbiota may be more successful at invasions (61,62).

## Conclusions

We demonstrate that host density is a major driver of the composition of microbiomes in *Durvillaea antarctica*. The majority of differences in microbiomes associated with bull kelp can be attributed to differences in beta-diversity associated with differing densities and tissue types of bull kelp. The greater beta-diversity found in low density populations is largely attributed to a stronger influence of dispersal limitation due to reduced intra-population interactions between hosts. Further work ought to examine the functional redundancy of these communities over large spatial scales, and study how these communities assemble immediately following recolonization. Finally, characterizing the sources of microbes in these communities is essential to understanding the recolonization process, and thus future work should aim to identify the degree to which vertical and horizontal transmission occurs, and how these interact with host dispersal.

## Acknowledgements

We thank Jordan Marsh of Clarence River Rafting for field support at Waipapa Bay and Mackenzie Dagg for her help with fieldwork, and to Jon Waters for helping to arrange some of this work. We also thank Doug Mackie and Linda Groenewegen for assistance in the lab. This research was supported by funding from the Royal Society of New Zealand (Rutherford Discovery Fellowship RDF-UOO1803 and Marsden Fund projects MFP-18-UOO-172 and MFP-20-UOO-173). WSP was supported by a University of Otago Doctoral Scholarship and Department of Marine Science PhD funding.

